# A bispecific antibody overcomes limitations of prior strategies to achieve effective and tumor-selective Death Receptor 5 (DR5) clustering

**DOI:** 10.1101/2025.01.10.632444

**Authors:** Victor S. Goldmacher, Iosif M. Gershteyn, Ravi Chari, Yelena Kovtun

## Abstract

The bispecific antibody IMV-M was designed to bind and cluster Death Receptor 5 (DR5) specifically upon engaging the tumor antigen MUC16. This dual-binding mechanism induces apoptosis in MUC16-positive tumor cells through a novel enhancement process. IMV-M exhibited potent, MUC16-selective anti-tumor activity *in vitro* and in xenograft models, achieving efficacy at safe therapeutic doses without requiring secondary crosslinking. In contrast, earlier DR5-targeting tumor-selective bispecific antibodies either demonstrated limited anti-tumor activity or relied on secondary crosslinking, highlighting that binding of a bispecific antibody to a tumor antigen and DR5 alone is insufficient for effective DR5 clustering. Our findings reveal that an additional crowding mechanism for efficient DR5 clustering by bispecific antibodies is required. In contrast to the targeted delivery approach with antibody-drug conjugates (ADCs), our approach offers a pathway for developing more effective and safer cancer therapies. Importantly, the MUC16 antigen is overexpressed in substantial subsets of ovarian, pancreatic, and lung cancers while being almost completely absent in normal tissues, with highly limited expression in certain epithelial cells, indicating broad applicability of this bispecific antibody.

## Introduction

With the advent of monoclonal antibodies, extensive research has focused on developing cancer therapies that target antigens expressed on the surface of cancer cells^1^. It soon became apparent that, to effectively eradicate tumors, most antibodies needed to be armed or possess an additional function^2^. Several strategies for enhancing antibody-based therapies have shown success, including antibody-drug conjugates (ADCs)^3^, Bispecific T-cell Engagers (BiTEs)^4^, and related CAR-T cell therapies^5^. However, these approaches demonstrate efficacy in only subsets of patients and are frequently associated with severe side effects. ADCs rely on endocytosis for payload delivery, which limits their ability to effectively deliver cytotoxic agents inside target cells, and their maximum tolerated dose is constrained by the inherent toxicity of the payload^3^. CAR-T and BiTE therapies, while potent, often trigger severe inflammatory responses^5,6^. Additionally, ADCs and CAR-T therapies require complex and expensive manufacturing processes. Consequently, there is an urgent need for new targeted therapies that offer improved efficacy, reduced toxicity, and greater cost efficiency.

As an alternative, another approach has emerged: bispecific apoptosis triggers (BATs). BATs are bispecific antibodies designed to target both tumor-associated antigens and death receptor 5 (DR5 or TNFRSF10B, TRAILR2)^7^. Apoptosis induced by DR5 occurs through the extrinsic apoptotic pathway. Upon binding its ligand TRAIL and clustering on the cell surface, DR5 recruits and activates pro-caspase-8 and/or pro-caspase-10^8^. These initiator caspases, once activated, cleave and activate downstream effector caspases, which ultimately leads to apoptotic cell death characterized by DNA fragmentation, membrane blebbing, and cell disassembly^8^. BATs are presumed to selectively accumulate on cancer cells, subsequently engaging and activating DR5 to induce apoptosis and eliminate tumors^7^. However, since effective signaling and apoptosis induction require DR5 clustering, monovalent or bivalent binding to DR5 is insufficient to induce apoptosis^9^. Early BAT development efforts were based on the premise that binding to an overexpressed tumor-associated antigen would create sufficient molecular crowding to induce DR5 clustering. However, the resulting BATs demonstrated limited efficacy in preclinical models and, not surprisingly, showed limited efficacy in clinical trials, indicating insufficient DR5 clustering by these agents^7^.

In this study, we introduce IMV-M, a bispecific antibody targeting the tumor-associated antigens MUC16 and DR5. IMV-M selectively activates DR5 on MUC16-positive tumors through a unique mechanism that uses the anti-MUC16 antibody component’s ability to bind multiple repeats in the MUC16 extracellular domain, bringing multiple IMV-M molecules adjacent to each other and thereby generating DR5 clustering.

## Results

### The bispecific antibody IMV-M was designed to effectively and selectively cluster DR5 on cancer cells

To generate a bispecific antibody that could strongly cluster DR5 on cancer cells, we took advantage of (i) the structure of MUC16, which, unlike the majority of cell surface antigens, contains multiple similar segments (repeats) in its extracellular domain^10^, and (ii) Sofituzumab (hu3A5), an anti-MUC16 antibody, which is capable of binding multiple adjacent antibody molecules to the repeats on the same MUC16 molecule^11^. We hypothesized that a bispecific antibody comprised of Sofituzumab fused to an anti-DR5 component could enforce effective DR5 clustering on MUC16-positive cells. To test this hypothesis, we created an antibody with the following structure (Fig. 1a): Sofituzumab IgG antibody was fused with a scFv fragment of a fully human anti-DR5 antibody via a flexible linker to allow an optimal geometry for DR5 clustering and activation. With this design, IMV-M was expected to promote DR5 clustering and subsequent activation on MUC16-positive cells (Fig. 1b), while on MUC16-negative cells, this bispecific antibody could bring together no more than two DR5 molecules (as shown in Fig. 1c). Furthermore, the high-affinity of this anti-MUC16 antibody (K_D_ ∼ 0.3 - 0.9 nM^12^) along with a low-affinity anti-DR5 scFv (K_D_ ∼ 0.2 µM, Supplementary Fig. 1, Supplementary Table 1), enhanced the selectivity of the construct’s interaction with MUC16-positive cancer cells.

**Fig. 1.**
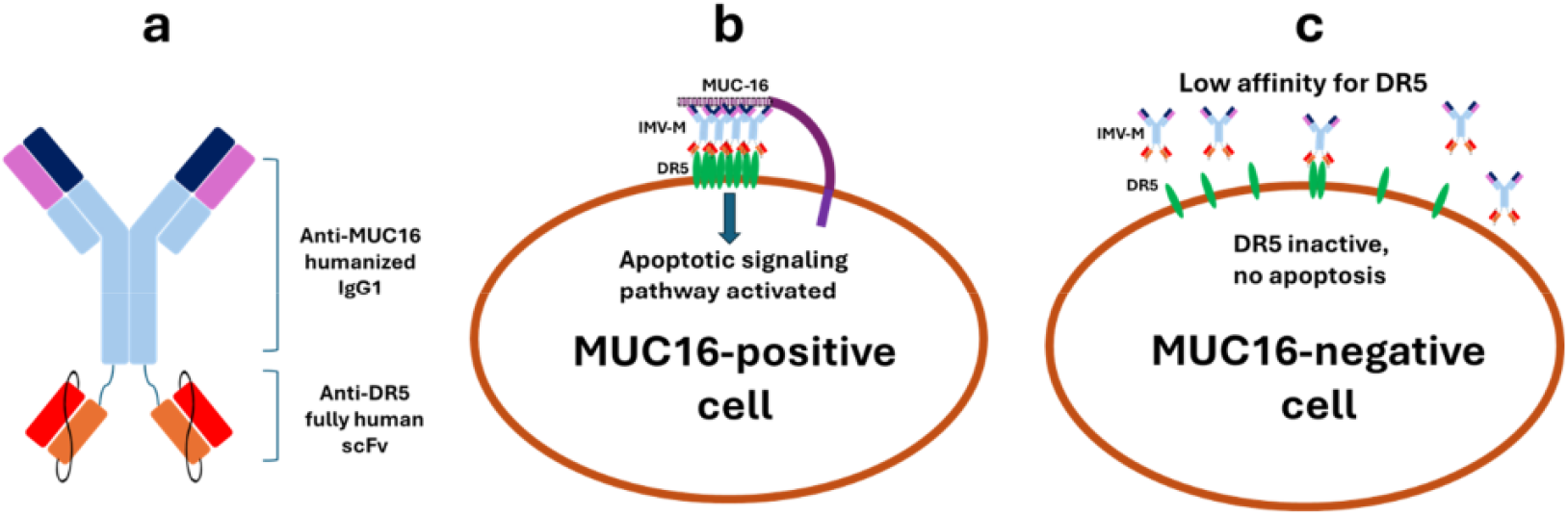
Design and Mode of Action of IMV-M. **a** IMV-M consists of the high-affinity anti-MUC16 humanized IgG1 antibody hu3A5, fused at its heavy chain C-terminus to a low-affinity scFv fragment of the anti-death receptor 5 (DR5) antibody lexatumumab, connected via a flexible linker. **b** Proposed mechanism of action of IMV-M: Multiple IMV-M molecules bind to the repetitive units of the cell surface MUC16 molecule, bringing them into close proximity to each other and the surface DR5 molecules. This proximity promotes the lateral diffusion of multiple DR5 molecules across the membrane, crowding them together. The flexible linker allows the DR5 molecules to align in an optimal geometric configuration, enabling clustering. Once DR5 clustering occurs, extrinsic apoptotic pathway is activated, and the cell undergoes programmed death and elimination. **c** IMV-M is designed to exhibit low affinity for DR5 to minimize binding to DR5 on the surface of MUC16-negative cells. Even in the rare instance where the binding occurs, the IMV-M molecules cannot cluster more than two DR5 molecules. These design features ensure that IMV-M cannot induce DR5 clustering or apoptosis in MUC16-negative cells.

### IMV-M inhibits the proliferation and induces apoptosis of MUC16-positive cells

We evaluated the cytotoxic activity of IMV-M after a two-day exposure in a panel of human MUC16-positive cell lines derived from pancreatic, breast, gastric, ovarian, and non-small cell lung cancers. As selectivity controls, we used the parental anti-MUC16 monospecific antibody and a bispecific antibody, consisting of an anti-fluorescein IgG^13^ fused to the same anti-DR5 scFv as used in IMV-M (Fig. 2a). Exposure of cells to IMV-M resulted in effective cell killing even at the lowest tested concentration (0.16 nM), whereas the anti-MUC16 antibody showed no cytotoxic effect across the entire tested concentration range (0.16–10 nM), and the anti-fluorescein/anti-DR5 bispecific antibody was unable to eradicate the cell populations even at the highest concentration (10 nM). We then evaluated the anti-proliferative and apoptosis-inducing activity of IMV-M in MUC16-positive PK-59 pancreatic adenocarcinoma cell populations in real-time monitoring cell proliferation and apoptosis progression to its final stages in individual cells. Exposure of the cells to IMV-M resulted in a nearly complete arrest of cell proliferation even at the lowest tested concentration (40 pM) within the first 24 hours (Fig. 2b) and activation of the effector caspases in most cells within the first 8 hours (Fig. 2c). In contrast, the anti-MUC16 antibody showed no effect on cell growth, as cell proliferation remained exponential, and the antibody did not induce apoptosis across the entire range of tested concentrations (40 pM to 10 nM). The anti-fluorescein/anti-DR5 antibody caused only a partial slow-down in cell proliferation and induced apoptosis in a small fraction of PK-59 cells, even at the highest tested concentration (10 nM). These results demonstrate that IMV-M is highly potent and selective in killing MUC16-positive cells via apoptosis.

**Fig. 2.**
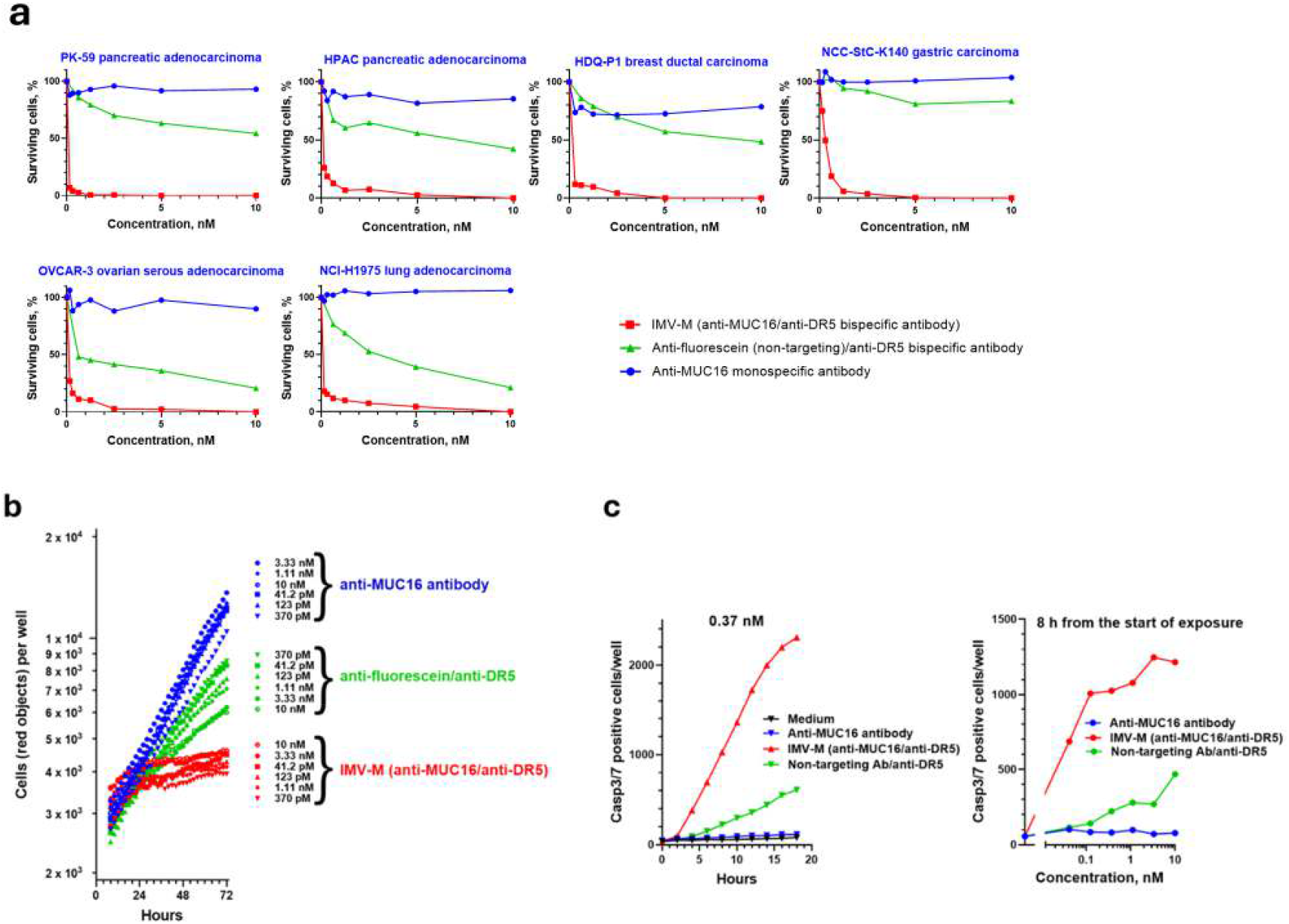
Cytotoxic effects of IMV-M on MUC16-positive cell lines in vitro. **a** IMV-M is selectively cytotoxic to MUC16-positive cell lines derived from pancreatic, breast, gastric, ovarian and non-small cell lung cancers. Cells were treated with various concentrations of IMV-M (red), a bispecific antibody of a similar design incorporating a non-targeting IgG1 and the same anti-DR5 scFv (green), or the monospecific parental anti-MUC16 IgG1 antibody (blue) for 2 days, and the relative number of viable cells was assessed using the CellTiter-Glo assay. **b, c** IMV-M arrests cell proliferation and induces apoptosis in PK-59 cells. Cells were treated with the same antibody conditions as in **a**, and individual cell proliferation, **b** and apoptosis, **c** were monitored in real time using the IncuCyte system as described in Methods. A representative subset of the apoptosis monitoring data is shown in panel **c**. Similar results were observed across the entire concentration range and throughout all monitored time points.

### IMV-M exhibits strong anti-tumor activity in several MUC16-positive xenograft models

To identify xenograft cancer models expressing MUC16, we used immunohistochemistry. NIH:OVCAR-3 xenografts served as a benchmark, as Genentech’s ADCs using the same anti-MUC16 antibody were previously tested on NIH:OVCAR-3 xenografts. As shown in Fig. 3 and summarized in Table 1, xenografts derived from various cancer cell lines displayed differing levels of MUC16 expression, ranging from strong (NIH:OVCAR-3) to negative (negligible). Our findings of MUC16 expression closely aligned with data from the Cancer Dependency Map Project at the Broad Institute (https://depmap.org/portal/). We then examined the anti-tumor activity of IMV-M in these xenograft models. In mice bearing established (∼ 130 mm^3^) xenografts of the pancreatic adenocarcinoma (PDAC)-derived cell line HPAC, a single intravenous injection of 5 mg/kg IMV-M resulted in pronounced tumor regression, while treatment with either a bispecific antibody containing a non-targeting IgG and the same anti-DR5 component, or with the parental anti-MUC16 monospecific antibody, had no impact on xenograft growth, similar to vehicle control (Fig. 4a). Furthermore, no body weight loss or any gross clinical abnormalities were observed in any animal group during the treatment period, indicating a lack of toxicity from IMV-M in mice (Supplementary Fig. 2). We investigated the dose-dependent anti-tumor activity of IMV-M using a xenograft model derived from the PDAC cell line PK-59. IMV-M demonstrated anti-tumor activity even at the lowest tested single dose of 1 mg/kg, with progressively stronger effects observed at the higher doses of 2.5 mg/kg and 5 mg/kg (Fig. 4b). Similar experiments with treatment at 5 mg/kg were carried out with several other PDAC and non-small cell lung cancer (NSCLC) xenograft models (Fig. 4C-G). IMV-M demonstrated strong anti-tumor activity in three out of five models with moderate MUC16 expression (HPAC, PK-59 and HCC827) and one out of two models with weak MUC16 expression (NCI-H1975). It showed modest activity in two models with moderate MUC16 expression (Capan-2 and SW1990) and one with low MUC16 expression (NCI-H292).

**Fig. 3.**
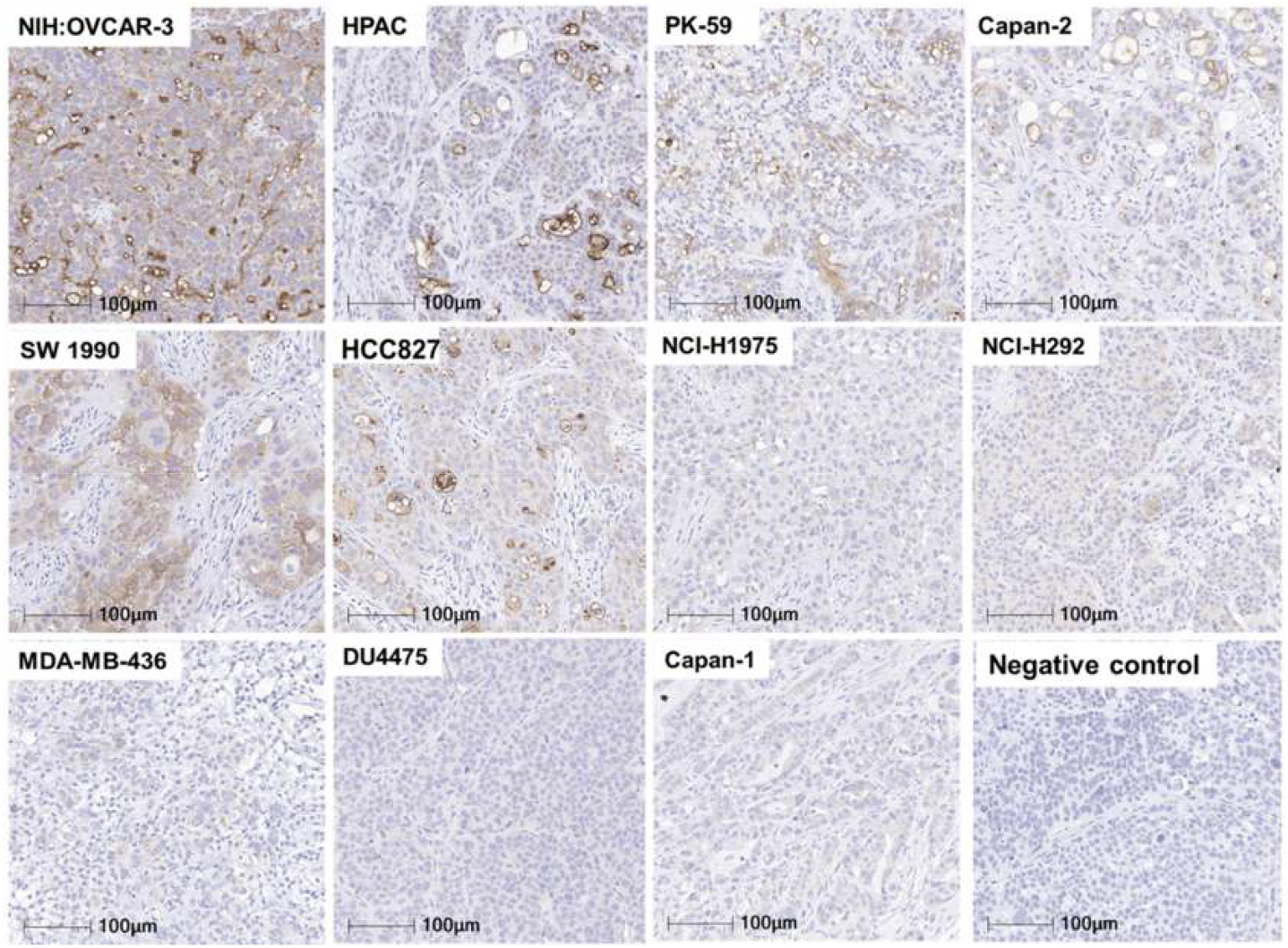
MUC16 Expression in Xenografts Derived from Cancer Cell Lines. MUC16 protein levels in xenograft tumors generated from cancer cell lines were evaluated using immunohistochemistry with an anti-MUC16 antibody. The summarized results are provided in Table 1.

**Table 1.** MUC16 Protein Levels in Tumor Xenografts (IHC Fig. 3)

| **Cell line** | **Origin** | **MUC16 expression** |
| --- | --- | --- |
| NIH:OVCAR-3 | Ovarian adenocarcinoma | Strong |
| HPAC | PDAC | Moderate |
| PK-59 | PDAC | Moderate |
| Capan-2 | PDAC | Moderate |
| SW1990 | PDAC | Moderate |
| HCC827 | NSCLC | Moderate |
| NCI-H1975 | NSCLC | Weak |
| NCI-H292 | NSCLC | Weak |
| MDA-MB-436 | Breast carcinoma | Negative |
| DU4475 | Breast carcinoma | Negative |
| Capan-1 | PDAC | Negative |

**Fig. 4.**
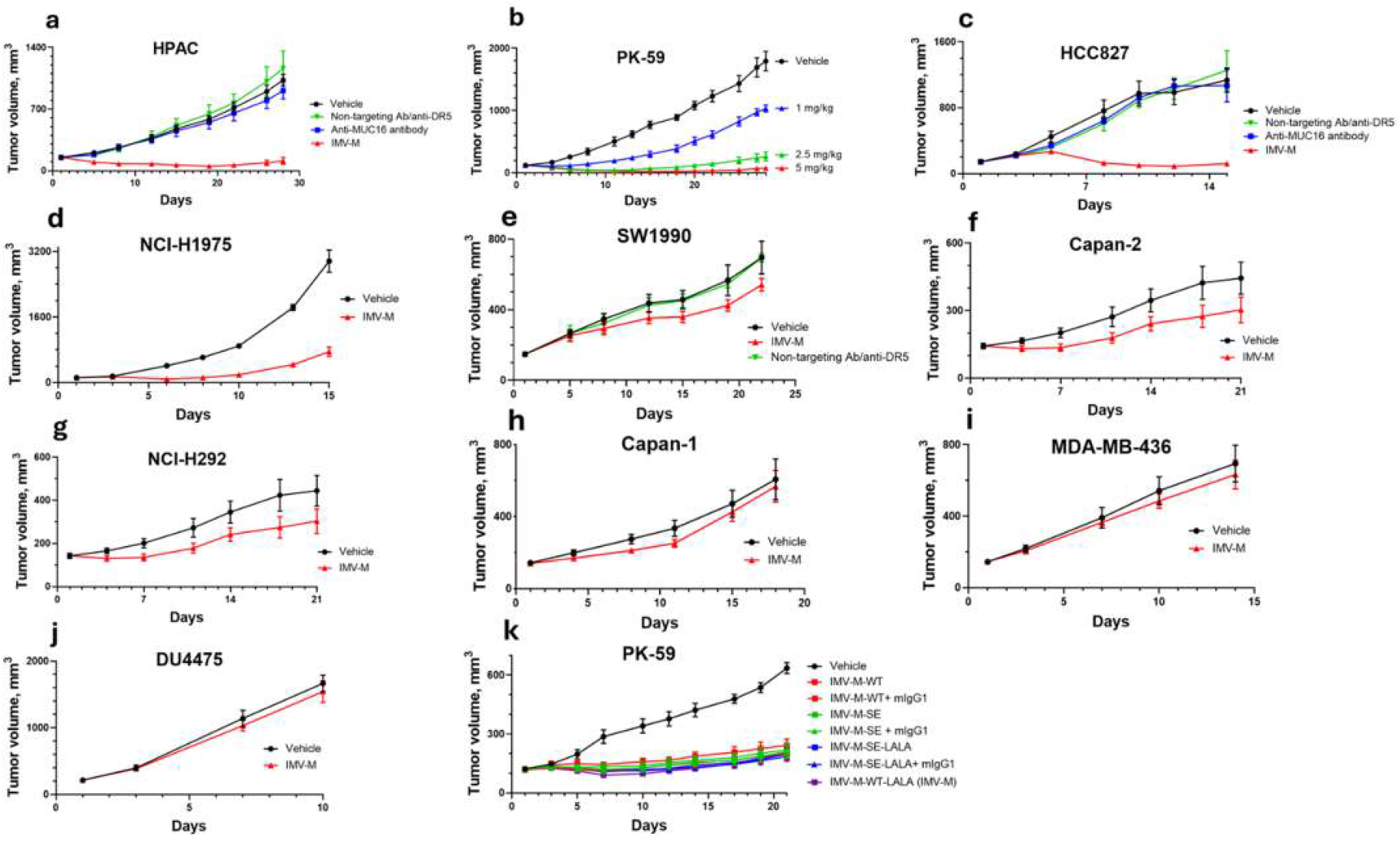
Anti-tumor activity of IMV-M in vivo in tumor xenograft models. **a** Nude mouse bearing established MUC16-positive HPAC PDAC-derived xenografts (∼130 mm^3^) were divided into four cohorts (n = 5 mice per group). The cohorts were treated intravenously on day 1 with IMV-M (red), a bispecific antibody of similar design incorporating a non-targeting IgG1 and the same anti-DR5 scFv (green), the monospecific parental anti-MUC16 IgG1 antibody (blue), or vehicle only (black), each at a dose of 5 mg/kg. **b** Nude mice bearing established MUC16-positive PK-59 PDAC-derived xenografts (∼130 mm^3^) were treated intravenously on day 1 (n = 4 mice per group) with IMV-M at 1 mg/kg (blue), 2.5 mg/kg (green), 5 mg/kg (red), or vehicle only (black). **c-g** Additional experiments were conducted using the same design as in **a** with MUC16-positive xenografts: HCC827 (n = 5 mice per group), NCI-H1975 (n = 5 mice per group), SW1990 (n = 5 mice per group), Capan-2 (n = 4 mice per group), and NCI-H292 (n = 4 mice per group). **h-j** Similar experiments were carried out with MUC16-marginal xenografts: Capan-1 (n = 5 mice per group), MDA-MB-436 (n = 4 mice per group), and DU4475 (n = 4 mice per group). **k** A separate experiment with MUC16-positive PK-59 xenografts tested variants of IMV-M containing Fc region mutations designed to enhance or reduce its affinity for FcγRII (n = 4 mice per group). Some groups were co-injected with 30 mg/kg of a non-targeting (anti-fluorescein) mouse IgG1. Data are presented as means ± SEM in all plots.

### IMV-M does not affect MUC16-negative xenograft models

To evaluate whether IMV-M may harm MUC16-negative tissues we conducted in vivo studies on xenografts with marginal or no MUC16 expression, based on immunohistochemistry (IHC) results (Fig. 3) and data from the Cancer Dependency Map. IMV-M demonstrated no adverse effects on these xenografts (Fig. 4H-J), reinforcing the likelihood that it would not be toxic to MUC16-negative normal tissues.

### The anti-tumor activity of IMV-M does not require interaction with the Fc gamma receptor II (FcγRII)

The *in vivo* anti-tumor activity of previously reported DR5-targeting monospecific IgG antibodies and the anti-FOLR1/anti-DR5 bispecific antibody depended on secondary crosslinking through interaction with FcγRII, which is expressed on B cells present in nude mice^14-16^. In the absence of FcγRII interaction, anti-DR5 antibodies lost their anti-tumor activity. This limited capacity of IgG1 anti-DR5 antibodies to cluster DR5 is attributed to their bivalency, restricting them to linking only two DR5 molecules at a time^7^. To assess whether the anti-tumor activity of IMV-M depended on FcγRII interaction, we compared the anti-tumor activities of several IMV-M variants at a 5 mg/kg dose:

1. IMV-M-WT-LALA: Featuring two mutations (L234A and L235A) in the human IgG1 Fc region, which significantly reduce IgG1’s affinity for FcγRII and other Fcγ receptors^17-20^.
2. IMV-M-SE-LALA: Incorporating the L234A and L235A mutations along with an S267E substitution, which selectively restores the affinity of human IgG1 for FcγRII^16,21^.
3. IMV-M-SE: Containing only the S267E substitution.
4. IMV-M-WT: Featuring no Fc-region mutations.

To simulate the high IgG1 concentration in human patients, which competes with therapeutic antibodies for Fcγ receptor binding, certain groups of mice were co-injected with 30 mg/kg of non-targeting mouse IgG1, achieving an approximate blood concentration of 2 µM. Notably, mouse IgG1 has a higher affinity for mouse FcγRII (KD of 0.3 µM) compared to human IgG1 (KD of 8 µM)^21,22^. As shown in Fig. 4K, the anti-tumor activity of IMV-M was unaffected by either enhanced or attenuated affinity for FcγRII or by the presence or absence of mouse IgG1.

### IMV-M exhibits favorable safety and pharmacokinetics in cynomolgus monkeys

Both the anti-MUC16 and anti-DR5 components of IMV-M cross-react with the corresponding cynomolgus monkey antigens^11,23,24^, indicating that the cynomolgus monkey species is a suitable preclinical model for evaluating the systemic toxicity of IMV-M. In a pilot study, two cynomolgus monkeys were intravenously injected twice with IMV-M, with a 21-day interval between doses; one monkey received 10 mg/kg per dose, and the other received 20 mg/kg per dose. Toxicity was assessed by monitoring hematologic parameters and blood clinical chemistry (Supplementary Table 2). No abnormalities were observed, indicating an absence of toxicity in this study.

The half-life of IMV-M in the blood of monkeys was also assessed (Supplementary Data). IMV-M circulation half-life was approximately 5-7 days and remained consistent after the second injection. These findings suggest that IMV-M may be sustained in the bloodstream long enough to effectively reach tumors, with no observed immune response leading to its neutralization in monkeys.

## Discussion

DR5 is an attractive target for inducing apoptosis in tumor cells, as it is broadly expressed across various cancers at levels sufficient to trigger apoptosis upon clustering and is upregulated in tumors relative to normal tissues^7^. Previously, anti-death receptor agents, including TRAIL derivatives and anti-death receptor antibodies, had achieved only limited success in both preclinical and clinical studies. Agents with minimal preclinical activity demonstrated negligible clinical efficacy, while those showing moderate preclinical effects similarly yielded only modest outcomes in clinical trials. No agents demonstrated consistently strong and broad preclinical efficacy unless enhanced by secondary crosslinking, resulting in an absence of substantial clinical impact (reviewed in^7^). This limited success likely arose from these agents’ inability to fully induce apoptosis. IMV-M stands as the first death receptor-targeting agent to exhibit compelling and broad anti-tumor activity in preclinical mouse models, indicating strong potential for clinical effectiveness.

MUC16 is a promising therapeutic target, as it is overexpressed in the tumors of most ovarian cancer patients and in a significant proportion of patients with pancreatic adenocarcinoma (PDAC) and non-small cell lung cancer (NSCLC). It is also present in subsets of patients with other cancers, including endometrial, breast, esophageal, gastric, and colorectal adenocarcinomas^25-37^. MUC16 is largely absent in most normal tissues, appearing only on the free surface of select epithelia^38,39^ [http://gepia.cancer-pku.cn; www.humanproteomemap.org; www.proteinatlas.org], which minimizes its exposure to circulating IMV-M.

Additional evidence supporting IMV-M’s clinical promise comes from studies with two ADCs, DMUC5754A and DMU46064A, which utilize the same anti-MUC16 antibody, hu3A5. Preclinical evaluations of these conjugates showed activity primarily in xenograft models with very high MUC16 expression, such as OVCAR-3 or greater^11,12^, and only at doses close to or exceeding their maximal tolerated doses in patients. Despite limited preclinical efficacy, these ADCs— especially DMU46064A—demonstrated promising activity (but narrow therapeutic window due to payload-related toxicity) in early clinical trials^40,41^. Given IMV-M’s robust efficacy in xenograft models with MUC16 expression levels notably lower than NIH:OVCAR-3 (Fig. 3) at doses several-fold lower than the predicted tolerable dose, it is likely to exhibit favorable clinical activity.

IMV-M offers a more effective and safer alternative to MUC16-targeting ADCs: (i) IMV-M acts directly on the cell surface, while ADCs require internalization and lysosomal degradation to activate. Given MUC16’s low internalization efficiency, ADCs need very high MUC16 expression levels to be effective, whereas IMV-M demonstrated activity even in xenografts with moderate MUC16 expression. (ii) IMV-M’s cell-killing mechanism is independent of drug resistance often acquired in chemotherapy-pretreated patients, making its activity less likely to be impacted by such resistance. In contrast, ADCs may be less effective in drug-resistant tumors. (iii) The maximum tolerated dose of ADCs is limited by the inherent toxicity of their payload, leading to a narrow therapeutic window, as seen with DMUC5754A and DMU46064A. Other types of MUC16-targeting therapies have shown limited success to date. For example, clinical trials of MUC16-targeted CAR-T therapies and a BiTE (Ubamatamab) have reported disappointing results^42,43^. There is substantial evidence suggesting that IMV-M will be safe for patients: (i) No signs of toxicity were observed in cynomolgus monkeys. (ii) The DR5 component within IMV-M has been previously incorporated in another bispecific antibody, and also is a fragment of the antibody Lexatumumab, both of which were well tolerated by patients in clinical trials^44-46^. (iii) No MUC16-related toxicity was observed in clinical trials of the two MUC16-targeted ADCs, and the systemic toxicity observed was consistent with the typical profile of ADCs using this cytotoxic payload^40,41^. Overall, IMV-M demonstrates superior preclinical efficacy compared to prior DR5- or MUC16-targeting agents, providing enhanced activity and safety and paving the way for more effective and safer cancer therapies.

## Methods

### Cell Culture

HPAC (CRL-2119), NIH:OVCAR-3 (HTB-161), NCI-H1975, SW1990, NCI-H292, Capan-1, Capan-2, DU4475, MDA-MB-436, NCI-H1975, and HCC827 were obtained from the American Type Culture Collection (ATCC). PK-59 was obtained from Cobioer Biosciences Co., Ltd and the RIKEN BioResource Research Center. NCC-StC-K140 was obtained from RIKEN, and HDQ-P1 was obtained from The Leibniz Institute DSMZ – German Collection of Microorganisms and Cell Cultures GmbH.

### Expression of IMV-M and control proteins

Monospecific and bispecific antibodies for this study were generated at WuXi Biologics and BioIntron using standard transient expression procedures in CHO cells, followed by isolation through protein A chromatography. When indicated, additional purification was performed using MSS-Superdex 200 chromatography. The integrity and purity of the antibodies were assessed using reduced and non-reduced SDS-PAGE and SEC-HPLC. Sequences of proteins generated in this study are reported in Supplemental data.

### In vitro cytotoxicity and apoptosis testing

The CellTiter-Glo tests (Promega Corporation) were performed at Pharmaron and HD Biosciences (WuXi AppTec). Cells were plated into flat-bottom tissue culture 96-well plates. The following day, test reagents were added for an additional two days. The relative number of viable cells was then assessed using the CellTiter-Glo assay, following the standard manufacturer’s protocol. Inhibition of cell proliferation and activation of apoptosis in PK-59 cells were examined by monitoring individual cells in real time using the Incucyte S3, a live-cell imaging and analysis system (Sartorius AG) at Pharmaron. Cells were plated into a 96-well flat tissue culture plate at a density of 1500 cells/well. The following day, test reagents and monitoring dyes were added to the cells, and fluorescent images of individual cells were captured every two hours following the manufacturer’s protocol. Incucyte Nuclight Rapid Red Dye for nuclear labeling (Sartorius Cat. No. 4717) and Incucyte® Caspase-3/7 Green Dye for Apoptosis (Sartorius Cat. No. 4440) were used to detect cell proliferation (number of red-fluorescent objects) and apoptosis (number of green-fluorescent objects with activated caspase-3 and/or 7) in accordance with the manufacturer’s protocol to monitor the execution phase of apoptosis.

### Immunohistochemistry (IHC) study

This study was performed at HD Biosciences (WuXi AppTec). Tumor tissues embedded in paraffin were cut with a microtome to a thickness of 4μm. After heat-induced antigen retrieval in Citrate 6.0 (MXB #MVS0066), sections were immersed in a 3% hydrogen peroxide solution for 5 minutes. To avoid nonspecific staining, sections were then incubated in Dako REAL™ Antibody Diluent (Code S2022) supplemented with 5% normal goat serum (Kangyuan #KY-01021) for 30 minutes at room temperature, followed by exposure to the anti-MUC16 antibody (Abcam #110640) at a 1:500 dilution in the same buffer overnight. The sections were then treated with HRP-conjugated Polyclonal Goat anti-Rabbit Immunoglobulins (Dako, cat# 4003) and visualized with diaminobenzidine according to the manufacturer’s protocol. The nuclei were stained with hematoxylin. Slides were scanned using an Aperio Scanner: Versa 8 (Leica) at 200X magnification. Images were opened with HALO software, and the pen tool was used to create an annotation layer, with necrotic areas excluded from the annotation layer.

### Mouse xenograft studies

These studies were conducted at Pharmaron and HD Biosciences (WuXi AppTec). All procedures related to animal handling, care, and treatment were performed according to guidelines approved by the Institutional Animal Care and Use Committee (IACUC) of HD Biosciences, adhering to the standards set by the Association for Assessment and Accreditation of Laboratory Animal Care (AAALAC). BALB/c nude female mice, aged 6-8 weeks at the time of inoculation and weighing 16-20 g, were used for the study. Tumor cells in the exponential growth phase were used for inoculation. Each mouse was inoculated subcutaneously on the right flank with tumor cells to establish tumor growth. Treatments were initiated when the mean tumor volume reached approximately 130 mm^3^. Mice were randomly assigned to treatment groups based on tumor volume to ensure similar average starting tumor sizes across groups. The test compounds were administered intravenously via the tail vein. Throughout the study, mice were monitored for tumor growth and general health, including behavior, mobility, food and water consumption, body weight, eye/hair condition, and other signs of abnormal effects.

### Exploratory Non-GLP Toxicokinetics and Safety Assessment in Non-Human Primates

Two naïve female cynomolgus monkeys were used in this study. Animal 1 received IMV-M via intravenous bolus administration at a dose of 10 mg/kg on Days 1 and 22, while Animal 2 received IMV-M via intravenous bolus administration at 20 mg/kg on the same days. Serum samples were collected at various time points, and IMV-M concentrations were measured using an enzyme-linked immunosorbent assay (ELISA), as described in the supplementary data. Hematology and clinical chemistry samples were collected at multiple time points. Hematology parameters analyzed included leukocyte count, red blood cell (RBC) distribution width, erythrocyte count, platelet count, hemoglobin, mean platelet volume, hematocrit, white blood cell (WBC) differential, mean corpuscular volume, mean corpuscular hemoglobin, reticulocyte count, and mean corpuscular hemoglobin concentration. Clinical chemistry parameters analyzed included alanine aminotransferase, creatinine, aspartate aminotransferase, calcium, total protein, phosphorus, albumin, total cholesterol, globulin, triglycerides, albumin/globulin ratio, total bilirubin, alkaline phosphatase, sodium, γ-glutamyltransferase, potassium, glucose, chloride, urea, and creatine kinase.

## Data availability

Derived data supporting the findings of this study are available from the corresponding author upon request.

## Acknowledgments

This work was supported by ImmuVia Inc

## Author contributions

V.S.G. and I.M.G. conceived and designed the study. V.S.G., I.M.G., R.C. and Y.K. provided scientific advice and analyzed the data. All authors contributed to the writing of the manuscript.

